# Initiation and Formation of Stereocilia during the Development of Mouse Cochlear Hair Cells

**DOI:** 10.1101/2024.03.23.586377

**Authors:** Suraj Ranganath Chakravarthy, Thomas S. van Zanten, Raj K Ladher

## Abstract

Stereocilia are apically located actin-protrusions found on the hair cells of the inner ear. At least three rows of stereocilia are arranged in a graded staircase pattern, which is vital for mechanosensation. Stereocilia form soon after the specification of hair cells. While these steps have been well-characterized in the avian auditory epithelium, the equivalent information in mice is lacking. Using scanning electron microscopy and super-resolution microscopy, we investigate stereocilia formation from hair cell specification stages in the mouse organ of Corti. Even before differentiation, we find that sensory progenitors, which will give rise to both hair cells and support cells, have a dense lawn of microvilli. Hair cell specialisation is first apparent as an enrichment in junctional actin, followed by the relocalisation of kinocilium into an eccentric position and the thickening of hair cell microvilli closest to the kinocilium. To determine actin signatures associated with hair cell development, we use a new analytical method to map cellular actin filament distribution during development. By nomalising relative actin filament density, we obtain insights into cuticular plate development and actin redistribution during the earliest phases of hair cell specialisation.

## INTRODUCTION

An end-point of the differentiation of hair cells is the generation of stereocilia. Stereocilia are the site of mechanotransduction in the inner ear, a function that is critically dependent on their structure (Furness et al., 1997). Each is made up of bundles of highly crosslinked parallel actin filaments (Tilney et al., 1989; Velez-Ortega and Frolenkov, 2019). Stereocilia are organised in graded rows arranged in a staircase pattern on the apex of the hair cell. These are placed next to an eccentric true cilium, the kinocilium. Together, this is called the hair bundle. The development of stereocilia has been well-characterized in avian systems (Tilney et al., 1992a; Tilney and DeRosier, 1986; Tilney et al., 1986) but in mammals, the events that mark the initiation of the stereocilia are unclear. Hair cells and their precursors are epithelial cells, and like most epithelial cells, hair cell precursors possess microvilli and a centrally located primary cilium. Microvilli, like stereocilia, are projections of actin filaments (Sharkova et al., 2023). In general, most epithelial cells, such as the intestinal enterocyte, have microvilli of similar length. In contrast, hair cells possess stereocilia that have varying length and width. Thus, even though stereocilia can be considered modified microvilli, it is the transition of microvilli to stereocilia that is a key event in the differentiation of a hair cell.

Stereocilia formation has been best characterised in the chicken basilar papilla (Tilney et al., 1992b), but has also been described in the mouse, rat and hamster cochlea (Kaltenbach et al., 1994; Lenoir et al., 1987; Li and Ruben, 1979; Zine and Romand, 1996). The initial steps of differentiation occur after the radial relocalisation of the kinocilium. Here, the microvilli of the nascent hair cell “bloom”. An elongation of the microvilli follows this before differential elongation of the stereocilia elaborates the staircase pattern. During differential elongation, extraneous shorter rows of microvilli are resorbed, giving rise to the final, mature, hair cell in the cochlea of most mammals. At each milestone, regulators of actin are likely involved in sculpting the microvilli to a staircase-patterned array of stereocilia (Avenarius et al., 2014). The effect of mutations of actin-regulating proteins, such as those that bundle, cap, sever or link actin to membranes as well as myosin motor proteins, have all been assessed to understand the proximate causes for the establishment and maintenance of the stereociliary bundle (Park and Bird, 2023). However, their effects on early developmental processes, before any overt specialisation of stereocilia, remains enigmatic.

Mutations in stereocilia proteins can show defects in other regions of the hair cell. All epithelial cells have a well-formed junctional collar of actin filaments that maintains integrity. In addition, stereocilia are inserted into a thick meshwork of cortical actin, analogous to the terminal web, known as the cuticular plate (Pollock and McDermott, 2015). Mutants in actin-regulatory proteins could show additional defect in these structures, confounding interpretations on stereociliogenesis. Thus, identifying signatures of actin throughout the hair cell during its development would provide comprehensive insights into the role of actin-regulators are various stages of stereocilia development. We characterise microvilli and stereocilia bundle development in the mouse inner ear, from embryonic to postnatal stages. Using a new analytical pipeline, we correlate morphological changes during stereocilia formation to patterns of actin localisation in the hair cell.

## METHODS

### Animals

CD1 mice were used for all experiments. Use of mice was carried out in accordance with institutional ethics guidelines.

### Immunohistochemistry

Dissection and fixation of inner ear in 4% paraformaldehyde (Sigma-Aldrich) overnight at 4⁰ C, was followed by the microdissection of the cochlea. For UN-RaPA analysis, samples using several protocols from different experiments were used. These differ by the type of permeabilisation and the blocking but are based on previously used protocols (Hirono et al., 2004; Singh et al., 2022). Phalloidin was added with the secondary antibody and incubated for between 40 minutes - 1 hour at room temperature. Both Alexa Fluor 488 and Alexa Fluor 555 conjugated Phalloidin (Thermo Fisher Scientific) were used. A polyclonal antibody to IFT88 (Proteintech) antibody was used at 1:500 dilution. A goat anti-rabbit secondary antibody conjugated with Alexa Fluor 546 (Thermo Fisher Scientific) was used at a 1: 500 dilution and incubated at room temperature for 1 hour. Slides were mounted with ProLong gold antifade (Thermo Fisher Scientfic) and imaged using an Olympus FV3000 (60x oil immersion objective of 1.42 NA) or Zeiss LSM 980 Airyscan 2 (63x oil immersion objective of 1.4 NA).

### Scanning electron microscopy

Preparation of samples for SEM were as previously described (Singh et al., 2022). Briefly, dissection of the inner ear in phosphate buffered saline and were then fixed in 2.5 % glutaraldehyde (Sigma-Aldrich) in 0.1M sodium cacodylate buffer pH 7.3 with 3 mM CaCl_2_ for at least 24 hours. For P11 and P12 stages, inner ears were dissected after decalcification with 120 mM ethylenediaminetetraacetic acid (EDTA) (Montgomery and Cox, 2016), although in our modified protocol, this is in 0.1M sodium cacodylate for 2 days This was followed by microdissection of the organ of Corti in 0.1M sodium cacodylate buffer. The samples were then post-fixed in 1% osmium tetroxide (Sigma-Aldrich) for 1 hour at room temperature. This is followed by treating the tissue with 0.5 % thiocarbohydrazide (Alfa Aesar). Another round of osmium tetroxide and thiocarbohydrazide treatment was carried out. After the last osmium tetroxide fixation, the sample was dehydrated through an alcohol series and were then subjected to critical point dehydration (Leica EM CPD300). The samples were mounted with double sided carbon adhesive tape and sputter coated (Coating thickness ∼10 nm). Then the samples were imaged in Zeiss Merlin Compact VP. SE2 mode was used in imaging with a voltage of either 5kV or 10 kV as indicated.

### Image Analysis

#### SEM

ImageJ was used to count microvilli/proto-stereocilia/stereocilia using the Cell-Counter plugin. We measured the widths of SEM images, of similar magnifications except for P11 and P12, by drawing a horizontal line across the shaft region of microvilli/proto-stereocilia/stereocilia and used the measure function (Hadi et al., 2020).

#### AiryScan

Image data was obtained from different experiments. The imaging settings were similar, but we ensured that saturation was avoided by changing laser settings. The default AiryScan processing option was used to obtain the images. We drew a horizontal line through the middle of the cells and used the dynamic radial slice functions to obtain the radial projections. To measure the doming of hair cells, a horizontal line was drawn from just below the junctional contacts of hair cells and supporting cells. A separate vertical line was drawn perpendicular to the horizontal line. This vertical line was drawn only up to the hair cell surface or base of the stereocilia or microvilli. In the case of a flattened structure, a vertical line was drawn to include only the cell-cell junctions. The length of the vertical line was the measure of doming length.

#### Unbiased Normalised Radial Projection Analysis (UN-RaPA) for F-actin Analysis

Multiple images from a z-stack (300 nm slices) of a 100 µm region of the organ of Corti were collapsed in a single maximum projection image. Each pixel (80-100 nm) in this image contains information of relative actin density. From this image cell types were identified, had their cell boundaries outlined and the fonticulus specified. These landmarks enabled the intensity values of each cell to be reprojected from Cartesian (x,y) into polar (theta, r) coordinates. The reprojection involved extracting a 3-pixel averaged intensity profile that started at the center-of-mass of each cell and extended to the cell boundary. Subsequently, this profile was rescaled from 0 to 1, being the center of mass and the cell-cell boundary, respectively. Intensity profiles were recorded each 6 degrees until the cell profile recording reached full circle, i.e., 360 degrees. The intensity profile that traverses the center of the fonticulus was defined as the 0 degrees angle and placed in the middle of the map. All other intensity profiles were reorganized accordingly, from -180 to 180 degrees. To be able to include the full cell boundary each profile was extended 40% from the cell boundary. In this way cells of similar identity but varying shapes and sizes from multiple F-actin images could be combined into a single intensity heat map. Images at different developmental timepoints subsequently granted access to the temporal progress of the F-actin distribution in the three cell types (GER, IHC, and OHC). It is important to note that large alterations to the general elliptical shape of cells cause deformation that render comparison impossible. Therefore, this method is only applicable up to P4-P5 for OHC. After this stage, their shape starts deviating substantially from elliptical (Etournay et al., 2010). We could compare the different cells across images because each maximum projection was normalised to the inherent intensity from the GER cell boundaries. Moreover, the rescaled and reoriented mapping of various cells allowed us to extract the averaged intensity only from selected regions, here the apical region of the cell.

#### Statistics and fits per figure

The statistical data has been organized in function of data representation in the figures. Starting from Figure 1 B-C, the data presented was obtained from cell-cell boundaries (j) in fluorescent images from multiple animals (n): Apex section of the organ of Corti j=4175 (n=3), j=1624 (n=1), j=4218 (n=3), and j=4214 (n=2) for developmental timepoints E14.5, E15.5, E16.5 and E17.5, respectively; Mid section of the organ of Corti j=3734 (n=3), j=3890 (n=5), j=4513 (n=9), and j=2995 (n=3) for developmental timepoints E14.5, E15.5, E16.5 and E17.5, respectively; Base section of the organ of Corti j=6127 (n=3), j=3253 (n=3), j=4533 (n=2), and j=878 (n=1) for developmental timepoints E14.5, E15.5, E16.5 and E17.5, respectively.

**Figure 1.**
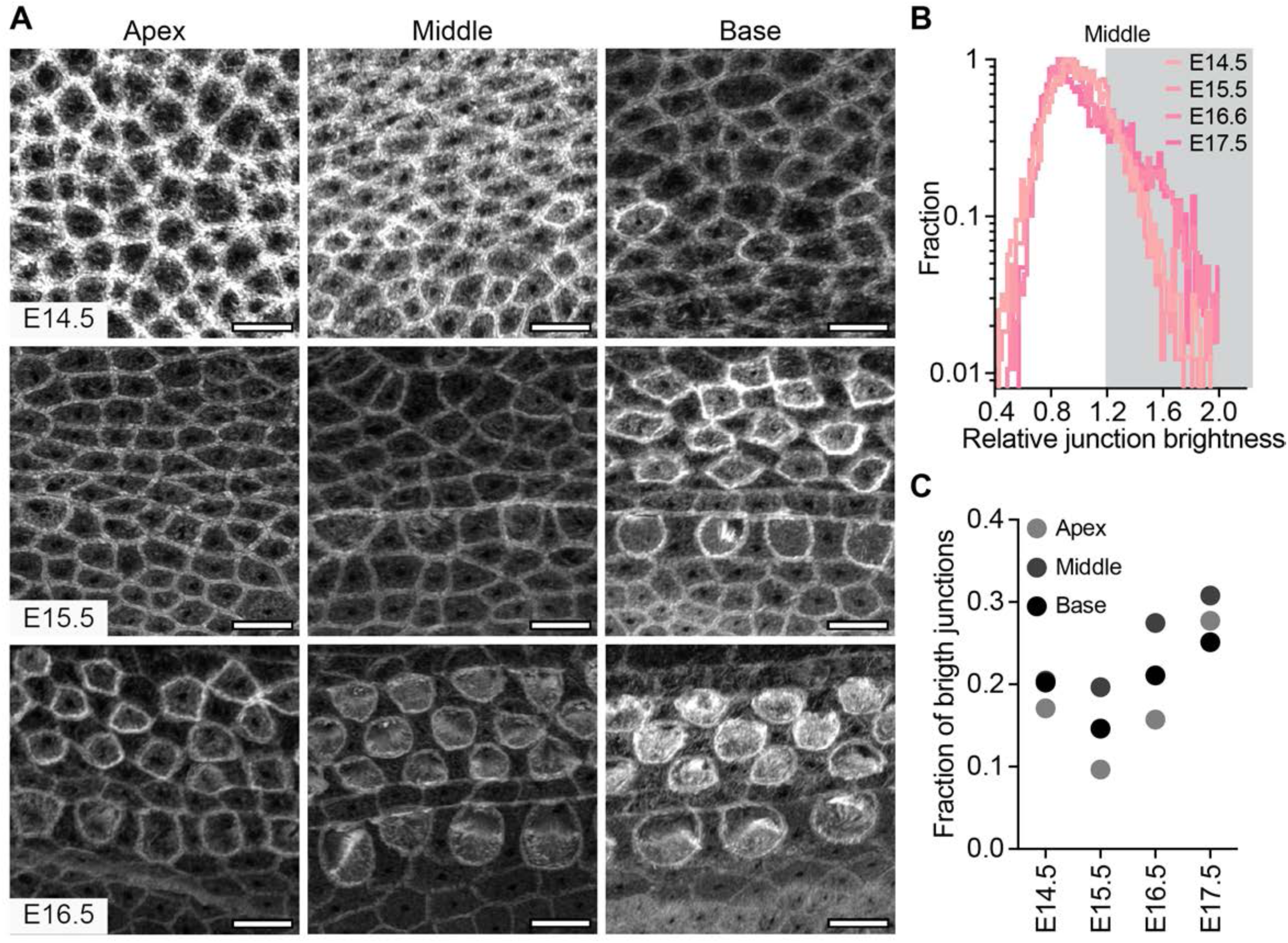
Differentiated hair cells have F-actin rich cell boundaries. (A) Super resolution airy scan images of phalloidin stained organ of Corti for the apical, middle, and basal region at E14.5, E15.5 and E16.5. Scale bar= 5 µm. (B) Logarithmic probability distribution of normalized cell-boundary intensity at different developmental timepoints at middle region of organ of Corti. (C) Fractions of cells exhibiting junctional intensities higher than 1.2 were plotted for apex, middle and base regions of organ of Corti at developmental timepoints E14.5 to E17.5.

For Figure 2B microvilli were counted from 3 (abneural) and 4 (neural) 8x8μm^2^ regions coming from different regions of the organ of Corti (n=2). To test for significance, we used a 2-tailed paired t-test (4 paired regions).

**Figure 2.**
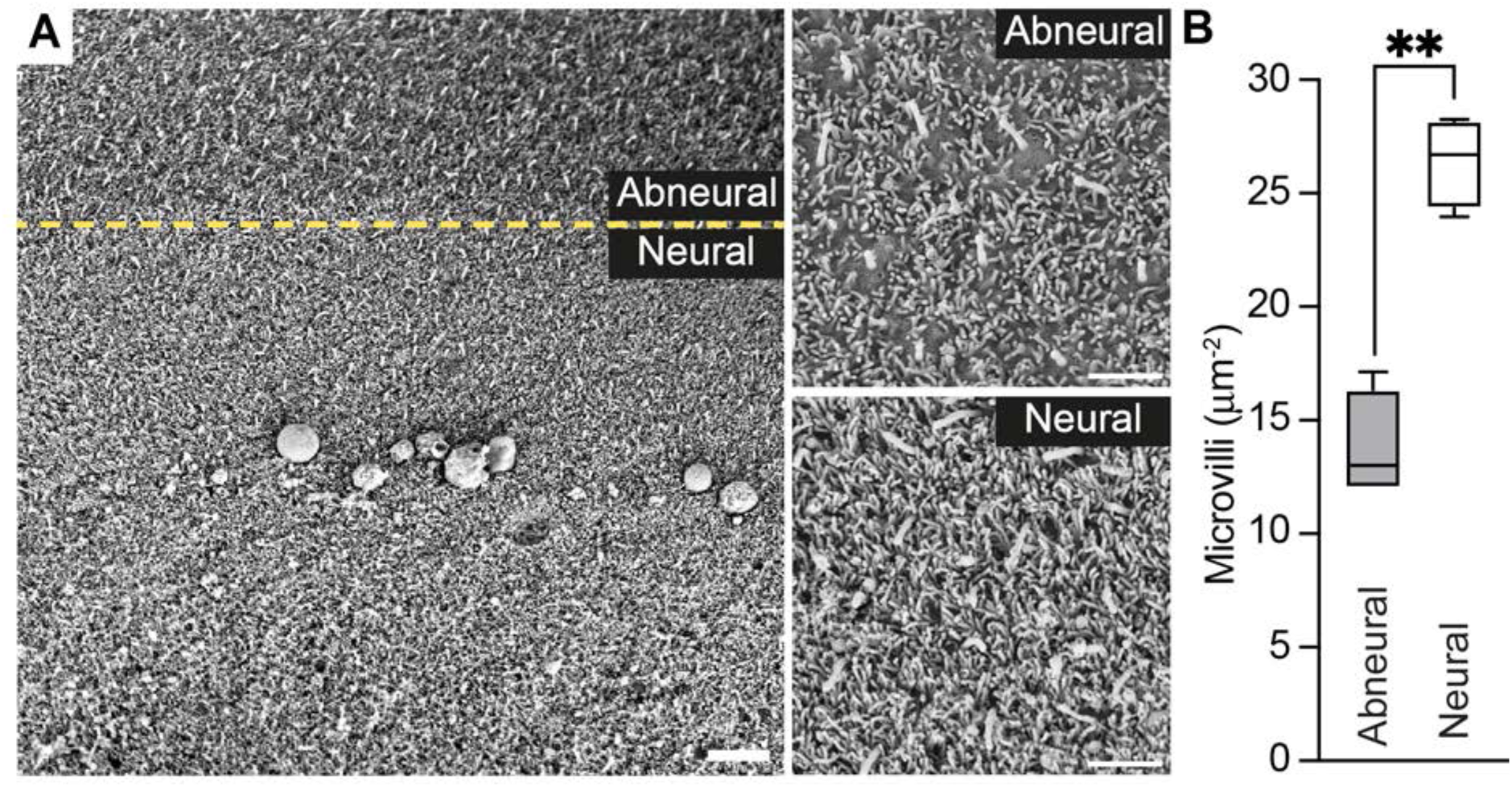
Density of microvilli in different regions organ of Corti. (A) To observe the density of microvilli in abneural and neural regions of sensory epithelium, large area scanning electron micrographs (SEM) of the organ of Corti were taken at E14.5. Scale bar = 5 μm. Dashed lines demarcate different regions annotated in the figure. Zoomed in regions of (A) display higher magnification SEM images showing microvilli at the surface of epithelial cells towards the abneural region and the neural region. The latter is the presumptive hair cell region of the organ of Corti at E14.5 (scale bar = 2 μm). All images are from the end of the middle region close to the apical turn of organ of Corti. (B) Quantification of the differences in microvilli coverage on abneural and neural regions, number of microvilli plotted per area (μm^2^). Microvilli were counted from 8x8μm^2^ regions, n=3 (abneural) and n=4 (neural) from 2 animals. Bars represent mean ± SD. ** p<0.005.

The heatmaps in Figure 3E are the average intensity of 36 OHC, 12 IHC, and 12 GER cells after reprojection.

**Figure 3.**
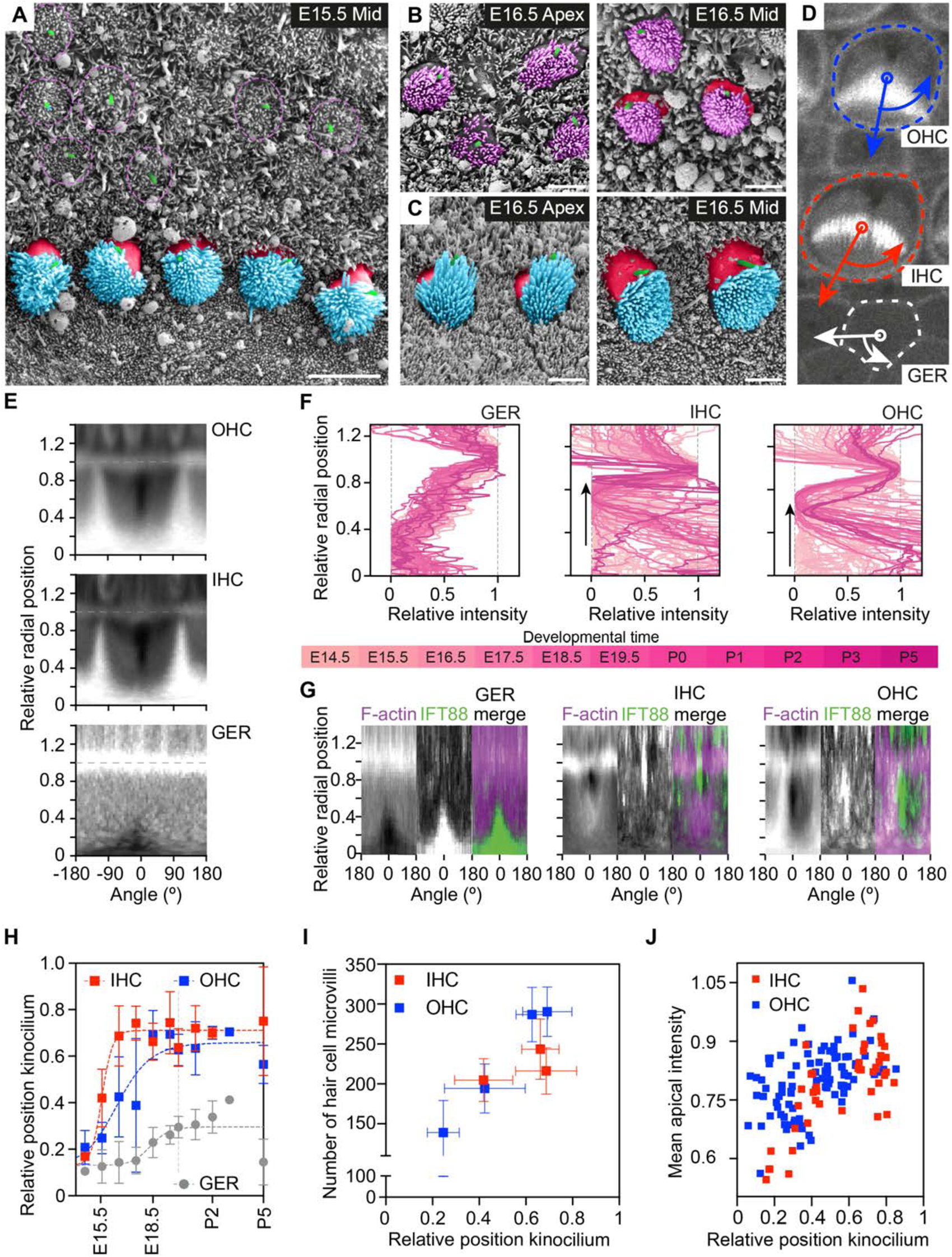
Relocalisation of kinocilia and initiation of Hair cell microvilli (HCmv) (A) SEM image of E15.5 organ of Corti. Scale bar = 5 μm. SEM image of OHCs (B panel) and IHC’s (C panel) at E16.5 apical and middle region of organ of Corti. Scale bar = 2 μm. False colouring of kinocilia (green colour), presumptive OHC (dotted purple circle), HCmv OHCs (purple), HCmv IHCs (cyan) and bare zone (red). Bilinear interpolation was used in the false colouring of the image in photoshop (D) Maximum projection confocal image showing an OHC, IHC and GER cell. For the UN-RaPA analysis, cells were identified, and their cell boundaries outlined (dashed line) together with their centroid (small circle). IHC were identified in red, OHC in blue and GER cells in white. The curved arrow marks the direction and larger straight arrow marks extent of the radial slice outside the boundary of cells. The organ of Corti was stained with phalloidin 488 and middle region of E19.5 is used. (E) The resulting UN-RaPA reprojection into (theta,r) showing an 2D heatmap for the three cell types. Each map is the average map from at least 10 cells. Dashed grey line at r=1 indicates cell boundary for the three heat maps. (F) Actin traces obtained at different developmental timepoints from a single radial slice along the 0° angle of GER cells, IHCs and OHCs. To facilitate visualization, each profile has been normalized so its minimal cellular value is set to 0 and the intensity at the cell-boundary corresponds to 1. (G) 2D heatmap of dual labelled organ of Corti stained with phalloidin and IFT88 showing the colocalisation between the actin-sparse fonticulus and the kinocilium. (H) Relative position of the fonticuli/kinocilia over developmental timepoints (from E15.5 to P5) tracks the relocalisation of kinocilium on hair cells and primary cilium on GER cells. (I) HCmv number and relative kinocilia position. The HCmv were counted using scanning electron microscope and the position of kinocilia was assessed using UN-RaPA from animals at similar developmental times (mid region of OC from E15.5 to P0). (J) Changes in the mean F-actin intensities on apical domain of HCs with kinocilium localisation (mid region of OC from E15.5 to E18.5). Note that each dot is a single cell.

Each line in Figure 3F corresponds to the 0-degree profile of the UN-RaPA projection image. Each of those images is derived from the average of at least 14 reprojected cells in an image coming from the indicated developmental timepoint. The number of images per timepoint were the following: E14.5 (n=3, 4 images), E15.5 (n=5, 11 images), E16.5 (n=8, 26 images), E17.5 (n=3, 7 images), E18.5 (n=5, 5 images). E19.5 (n=3, 6 images), P0 (n=4, 5 images), P1 (n=4, 14 images), P2 (n=3, 5 images), P3 (n=1, 1 image), and P5 (n=1, 4 images).

The heatmaps of Figure 3G correspond to the averaging of 15 GER, 21 IHC, and 68 OHC cells after reprojection. Both F-actin and IFT88 reprojection was performed from the same positions.

Figure 3H displays the mean±sd of the peak position of the Gaussian fit to each of the lines depicted earlier in Figure 3F. The cumulative number of cells per developmental timepoint of the different cell types are: E14.5 (30 GER, 30 IHC, 24 OHC), E15.5 (116 GER, 147 IHC, 297 OHC), E16.5 (314 GER, 383 IHC, 1221 OHC), E17.5 (66 GER, 85 IHC, 273 OHC), E18.5 (53 GER, 67 IHC, 225 OHC). E19.5 (67 GER, 78 IHC, 253 OHC), P0 (55 GER, 58 IHC, 200 OHC), P1 (109 GER, 158 IHC, 524 OHC), P2 (48 GER, 63 IHC, 193 OHC), P3 (14 GER, 16 IHC, 61 OHC), and P5 (33 GER, 48 IHC, 151 OHC). Each of the cell type datapoints have been fitted with a 4-parameter sigmoidal fit displayed in the graph as a dashed line.

The data in Figure 3I is a composite of counted microvilli data (also discussed below) and the UN-RaPA analysed fluorescent data mentioned above. The microvilli number (y-axis) in the three data points (mean±sd) belonging to IHC came from mv counted on 26 cells (E15.5; n=2), 20 cells (E16.5; n=2), and 15 cells (E18.5; n=3). The values on the x-axis belonging to these data points are the mean±sd from 147 cells (E15.5; n=5, 11 images), 383 cells (E16.5; n=8, 26 images), and 67 cell (E18.5; n=5, 5 images). For the OHC the microvilli number in the four data points (mean±sd) came from mv counted on 18 cells (E15.5; n=2), 23 cells (E16.5; n=2), 16 cells (E18.5; n=3), and 21 cells (P0; n=3). The values on the x-axis belonging to these data points are the mean±sd from 297 cells (E15.5; n=5, 11 images), 1221 cells (E16.5; n=8, 26 images), 225 cells (E18.5; n=5, 5 images), and 200 cell (P0; n=4, 5 images)

The data displayed in Figure 3J comes from 47 fitted IHC cells and 96 fitted OHC cells from developmental timepoints E15.5-E18.5.

Doming height measurements reported in Figure 4E were obtained from the following developmental timepoints: E14.5 (14 GER, 17 IHC, 17 OHC), E15.5 (29 GER, 10 IHC, 40 OHC), E16.5 (13 GER, 7 IHC, 27 OHC), E17.5 (9 IHC, 28 OHC), E19.5 (3 IHC, 8 OHC), P1 (6 GER, 7 IHC, 22 OHC).

**Figure 4.**
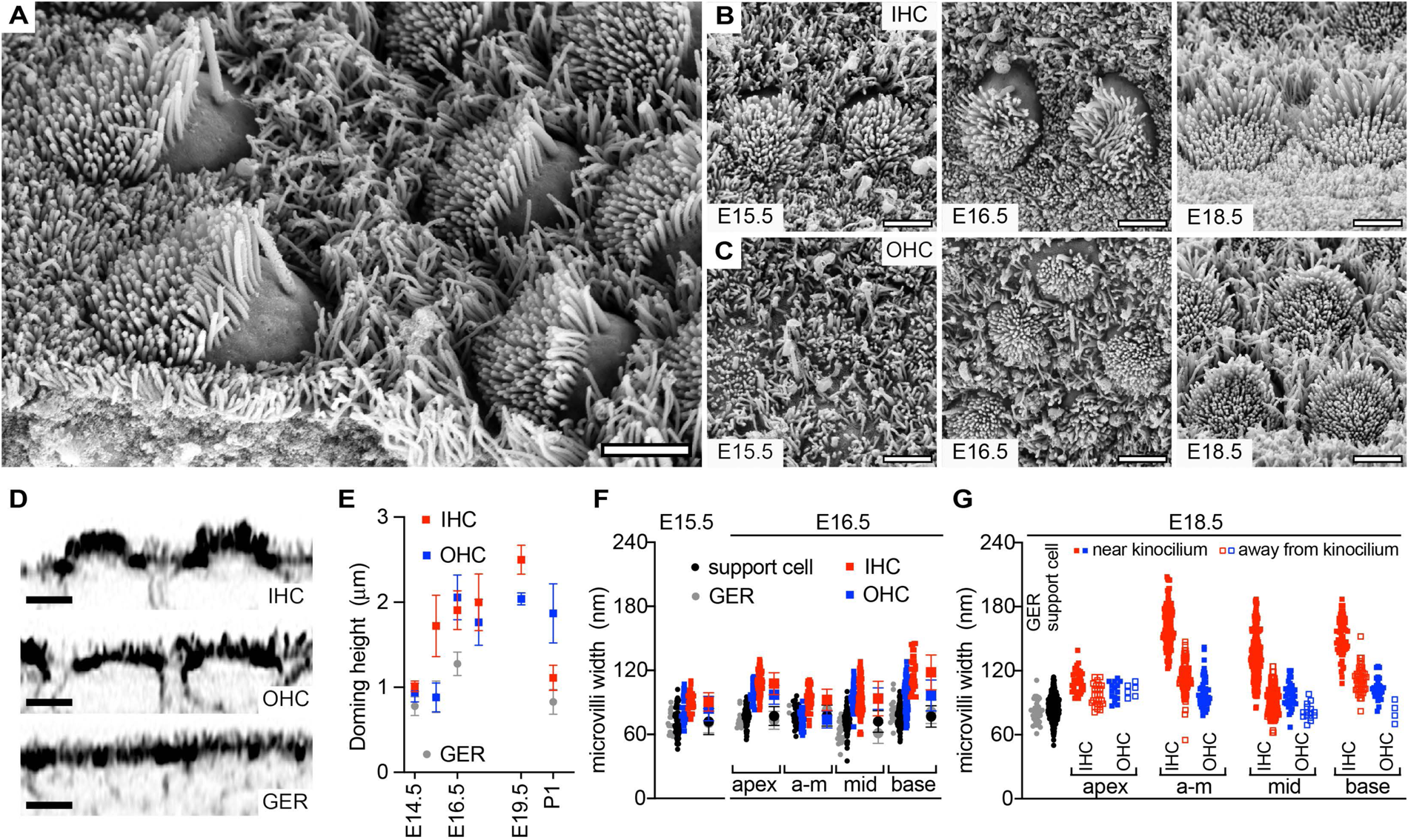
Sequence of events at embryonic stages leads to the specialisation of HCmv. (A) SEM (sideview) of the organ of Corti in basal turn at E18.5. Scale bar = 2 μm. SEM images of (B) IHCs and (C) OHCs at different developmental timepoints. All images are from the end of the middle region of organ of Corti. Scale bar = 2 μm. (D) x-z slices of airyscan images of organ of Corti at E15.5 base stained with phalloidin 488. Doming was assessed in IHCs, OHCs and GER cells. Scale bar = 2 μm. (E) Quantification of doming of apical surface in the IHC, OHC and GER cells across developmental timepoints from E14.5 to P1. The cells were in the basal region of organ of Corti. (F) Graphs with widths of HCmv on IHCs (red) and OHCs (blue), microvilli on support cells (SCmv) (black) and microvilli on greater epithelial cell (GERmv) (grey) at the middle region of the organ of Corti at E15.5 and apical, interface of middle and apical, middle and basal regions of the organ of Corti at E16.5. (G) Graphs displaying the widths of microvilli on support cells (SCmv: black), the widths of microvilli on greater epithelial cell (GERmv: grey) and the widths of HCmv on IHCs (red) and OHCs (blue), separated in two pools based on their proximity to kinocilia (closed or open symbols for near or away from the kinocilium, respectively). The data collates microvilli widths from the apical, interface of middle and apical, middle and basal regions of the organ of Corti at E18.5.

Microvilli width, Figure 4F, was obtained at E15.5 for IHC (31 mv, 6 cells, n=2), OHC (29 mv, 9 cells, n=2), GER cells (11 mv, n=1) and Support cell microvilli (69 mv, n=2). For the E16.5 data we obtained the following: apex IHC (60 mv, 11 cells, n=2), OHC (27 mv, 7 cells, n=1), GER cells (12 mv, n=1) and support cell (80 mv, n=2); mid-apex IHC (55 mv, 3 cells, n=1), OHC (51 mv, 7 cells, n=1), GER cells (2 mv, n=1) and support cell (26 mv, n=1); mid IHC (63 mv, 15 cells, n=2), OHC (57 mv, 19 cells, n=2), GER cells (20 mv, n=1) and support cell (97 mv, n=2); base IHC (31 mv, 8 cells, n=2), OHC (81 mv, 16 cells, n=2), GER cells (17 mv, n=1) and support cell (94 mv, n=2).

The E18.5 data in Figure 4G displays microvilli from GER cells (33 mv, n=1) and support cells (319 mv, n=2). Furthermore, a distinction is made between microvilli positioned near or away from the kinocilium. For the E18.5 data we obtained the following: apex IHC (36/25 mv, 5 cells, n=1), and OHC (15/0 mv, 4 cells, n=1); mid-apex IHC (136/106 mv, 7 cells, n=1), and OHC (16/30 mv, 4 cells, n=1); mid IHC (204/161 mv, 11 cells, n=2), and OHC (37/15 mv, 7 cells, n=2); base IHC (61/41 mv, 5 cells, n=2), and OHC (30/4 mv, 3 cells, n=1). For any data comparison to understand statistical significance an ordinary ANOVA test was performed.

The width measurements for Figure 5C-D were obtained for rows 1-3 on cells at the following developmental timepoints: E18.5 (204/0/0 on 11 IHC, n=2; 37/0/0 on 9 OHC, n=2), P0 (212/0/0 on 15 IHC, n=3; 108/0/0 on 8 OHC, n=2), P2 (79/52/81 on 15 IHC, n=3; 55/45/73 on 22 OHC, n=3), P4 (74/39/59 on 9 IHC, n=3; 19/18/49 on 11 OHC, n=2), P7 (43/23/39 on 8 IHC, n=2; 26/26/59 on 15 OHC, n=2), P11 (33/17/8 on 8 IHC, n=2; 10/11/28 on 2 OHC, n=1), and P12 (18/7/5 on 5 IHC, n=1). The (proto-)stereocilia increase in width was fitted to a linear curve for all three rows, for both the IHC and OHC.

**Figure 5.**
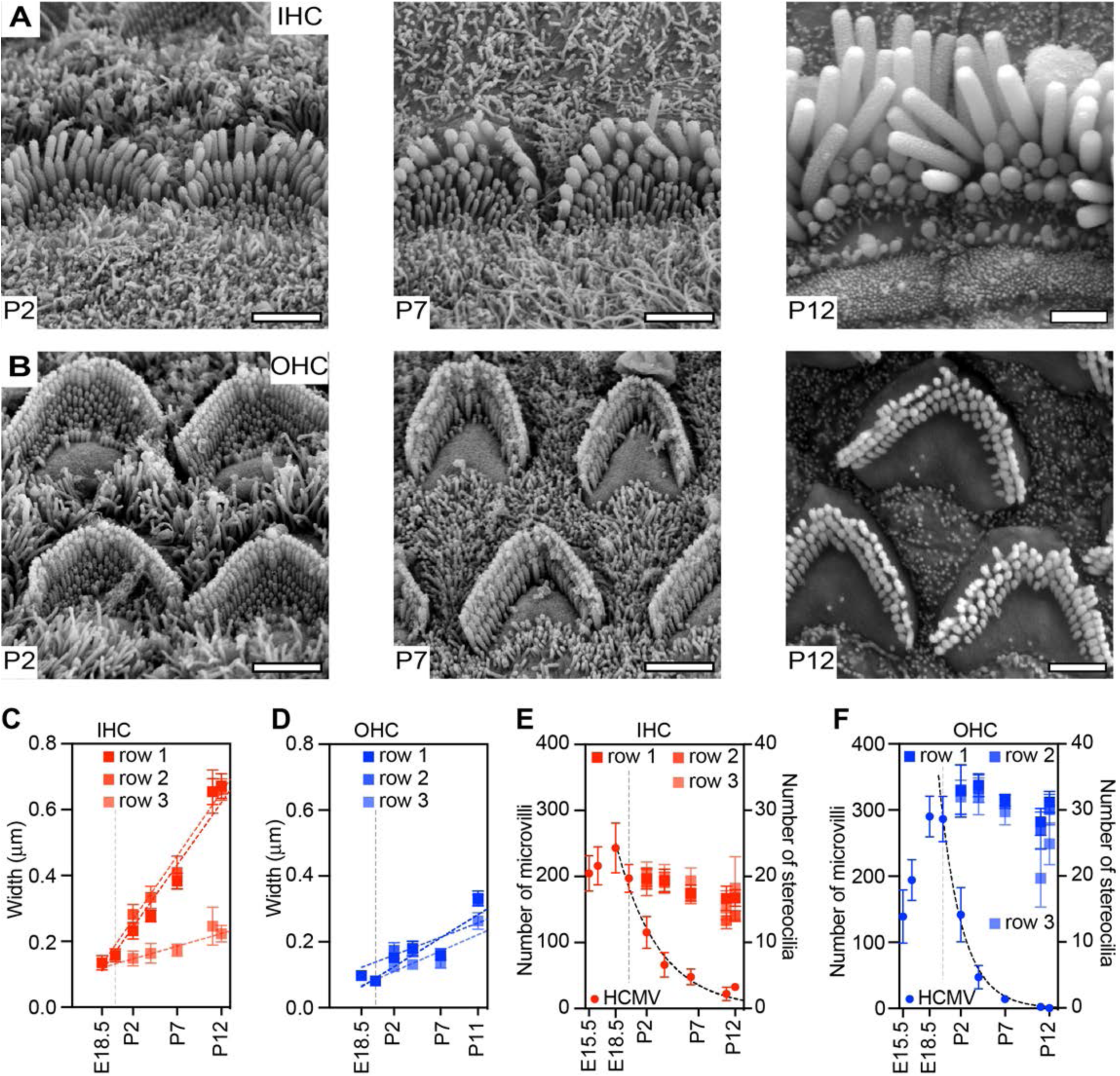
Development of Stereocilia. (A-B) Panel of SEM images at different developmental timepoints of IHCs (A) and OHCs (B). (C-D) Widths of the three rows of stereocilia with row 1 closest to kinocilia at different developmental timepoints on (C) IHCs and (D) OHCs. Number of HCmv, and stereocilia in row1, row2 and row 3 on (E) IHCs and (F) OHCs as a function of developmental time.

For Figure 5E-F the actin protrusion number per cell mean±sd were calculated from the following number of cells (and animals) per developmental time point: E15.5 (26 IHC, n=2; 18 OHC, n=2), E16.5 (20 IHC, n=2; 23 OHC, n=2), E18.5 (15 IHC, n=3; 16 OHC, n=2), P0 (19 IHC, n=3; 21 OHC, n=3), P2 (13 IHC, n=3; 18 OHC, n=3), P4 (15 IHC, n=3; 13 OHC, n=4), P7 (14 IHC, n=3; 16 OHC, n=3), P11 (6 IHC, n=2; 46 OHC, n=2), and P12 (4 IHC, n=1; 12 OHC, n=1). The microvilli number decrease was fitted to a single exponential decay (dashed curve) that started at E18.5 or E19.5, for IHC or OHC, respectively.

## RESULTS

### Enrichment of actin filaments in hair cell junctions during early differentiation

Hair cells become morphologically apparent, basally, at E18, with maturation progressing to the apical ends of organ of Corti (Chen and Segil, 1999; Li and Ruben, 1979; Lim and Anniko, 1985). The expression of Atoh1, which specifies HC, has a similar profile, although its expression is already detected in putative HC at E14.5, extending to the apex by E17.5 (Chen et al., 2002). To identify the first features of HC differentiation, we stained the organ of Corti from E14.5 to E16.5 mice with phalloidin (Figure 1A). We found that at E14.5, some cells at the base and the middle regions show enriched junctional actin. By E15.5, a greater number of cells with enriched junctional actin are observed in those regions, with some becoming detected in the apical regions as well. Cells that showed these enriched junctional actin belts never neighboured each other, and we hypothesised that these were presumptive hair cells. These cells showed a typical pattern of hair cell development, forming a chain-like pattern along the basal-apical axis of the organ of Corti, with presumptive inner hair cells appearing before presumptive outer hair cells (Cohen et al., 2023; Zine and Romand, 1996). By E16.5, these presumptive HC, identified by enriched junctional actin, were detected throughout the organ of Corti. In the basal regions, these cells also showed sub-cellular actin accumulation resembling nascent stereociliary bundles.

To quantify actin enrichment, we used tissue analyser to mark every cell junction in the field of view (Aigouy et al., 2016). The average intensity from each junction was measured and plotted as a frequency distribution for the basal, middle (Figure 1B) and apical regions of the organ of Corti from E14.5 to E17.5. A population of junctions showing higher F-actin accumulation could be detected starting from E14.5 and progressively increased with time as well as apically (Figure 1C). We also observe that the increase in the population of junctions with higher F-actin intensities at the apical region of OC correlated with the lag in the maturation of apical HCs. The similarity of the proportion of cells showing higher F-actin to the pattern of development of hair cells, suggests that the cells with higher accumulation of F-actin in their boundaries are likely to be hair cells. We further propose that this F-actin signature could be used to distinguish early differentiating hair cells from other cells in the organ of Corti.

### Microvillar Density defines Early Domains in the Organ of Corti

Although confocal microscopy showed differences in the levels of actin enrichment in clearly defined structures such as junctional cell boundaries, it is unable to resolve potential differences in features such as actin projections at the apical site of cells in the developing cochlea. To resolve these, we used scanning electron microscopy. While all cells showed microvilli (mv) we observed a significant difference in microvillar density at E14.5 between the abneural and neural regions of the organ of Corti (Figure 2A). A domain of low density mv was observed in lateral, abneural, regions of the organ of Corti (Figure 2B). In medial, neural regions, mv density was higher (Figure 2B). Nonetheless, the putative sensory patch cells that are present in the neural region, did not show any heterogeneity with their neighbours, despite our observation of enriched actin in junctions of some of the cells.

### Kinocilia Relocalisation correlates with increased HC microvilli number

The first morphological indication of HC specialisation was an increased microvilli number when the kinocilia radially relocated, a process that has been termed blooming. Consistent with this, SEM imaging showed that putative IHC in the middle region at E15.5 have increased mv number (Figure 3A-C). To ask if we could identify signatures of blooming we combined information from SEM with analysis from F-actin fluorescence confocal images. To compensate for cell-to-cell variability we reprojected the F-actin intensity values of each cell type into a normalized radial projection that will reorient and reshape each cell into a matching and unbiased cartograph. Moreover, to rectify any intensity variability between images due to variability in phalloidin staining or imaging conditions all intensity values were normalized to the average cell boundary intensity of the greater epithelial ridge (GER) cells. By determining the outline of each cell and pinpointing the F-actin sparse region (Figure 3D) the Unbiased and Normalised Radial Projection Analysis (UN-RaPA) presents an intensity map for each cell type (Figure 3E). In the F-actin intensity heat map, the cell boundary is clearly visible as the horizontal intensity region in the heat map at r=1 (Figure 3E; dashed gray lines). Intensity variation below this line displays the distribution of subcellular apical actin content. GER cells, for example, display very little intensity variability within their apical cell cortex aside from a local decrease in F-actin content. This actin-sparse region of the cell apex is called the fonticulus of an epithelial cell and is occupied by the cilium.

A summary of the intensity profiles, running from the center-of-mass until the cell-boundary at 0 degrees over developmental time, reveals limited profile change for GER cells (Figure 3F). In contrast, IHC or OHC cell types display dramatic profile changes. Dual color imaging with the ciliary marker IFT88 (Jones et al., 2008; Taulman et al., 2001), confirmed that F-actin fonticuli correspond to the cilium position (Figure 3G). We were, therefore, confident in using the position of the actin-sparse fonticulus region as a proxy for the position of the cilium/kinocilium at different stages. At each developmental time-point, multiple profiles (each an average of >10 cells per z-stack) were fit and combined to give an average ± standard deviation position relative towards the cell boundary (Figure 3H). Using this analysis, we observed that relocalisation of the kinocilium was first detected for putative IHC of the mid-base region of the OC at E15 and in the middle region by E15.5 (Figure 3H). Kinocilial relocalisation for OHC was around a day later, at E16.5 in the middle region (Figure 3H). The radial migration of kincolilia has been associated with the increase in mv number in putative hair cells (Tilney et al., 1992a; Zine and Romand, 1996). We therefore compared the measured number of mv obtained from SEM with the calculated kinocilium position obtained from UN-RaPA. This showed a positive correlation between mv numbers and kinocilial eccentricity for both IHC and OHC (Figure 3I).

We next asked whether signatures of microvilli blooming could also be observed directly in the fluorescence images of F-actin. While optical microscopy cannot resolve individual microvilli, the increased local density of microvilli should correspond to an increase in F-actin concentration and therefore an increased intensity. The advantage is that kinocilium position and mv density can be extracted from a single cell compared to tissue-averaged values. Scatter plots of IHC and OHC display a positive correlation between kinocilium displacement towards the cell boundary and the increase in F-actin density on the apical side of the hair cell (Figure 3J).

Altogether, both SEM and confocal data demonstrate clear signatures of mv blooming in developing mouse hair cells. Moreover, UN-RaPA data suggests that the tissue-level temporal developmental program is recapitulated in single cells.

### Hair Cells Dome during Differentiation

We observed that during differentiation, putative hair cells showed doming, where the apex becomes elevated in comparison to the surrounding support cells (Figure 4A). This is particularly prominent from E16.5 to E18.5 in both IHC and OHC (Figure 4B-C). Quantifying Airyscan images along the x,z plane reveals that IHC doming can be first detected at E15.5 (Figure 4D). Here doming is defined as the height difference between the most apical F-actin cortex and the cell-cell junctional F-actin accumulation. At E15.5 this height difference is 1.7±0.4 micrometer for IHC and reaches a maximum height of approximately 2.5±0.2 micrometer at E19.5 and flattens as the hair cell further develop (Figure 4E). In OHC, a doming of 2.1±0.3 micrometer is detected at E16.5 and appears to be oblate, with the region around the deflected kinocilia being the highest point of the dome (Figure 4A and 4E). Due to doming, we were unable to accurately measure the length of the microvilli, although they appear to show some extent of differentiation lengthening.

### Specialisation of Microvilli to Stereocilia

Doming of hair cells precluded quantitation of microvilli length. Instead, we asked if quantifying the microvilli width could provide insights into the specialisation of stereocilia. To better understand the pattern of Hair Cell microvilli (HCmv) thickening, we used the developmental gradient of HC differentiation (Figure 1A and Figure 4B-C). In the mid-apical region of OC at E15.5, we observe that the widths of HCmv on both presumptive IHCs and OHCs are equivalent (91±9 nm and 85±11 nm, respectively) and thicker (P<5·10^-4^) than the microvilli on support cells (73±12 nm) and GER cells (71±9 nm) (Figure 4F). This is the first evidence of heterogeneity amongst the microvilli in the sensory patch. IHCs in the mid-basal region bloom at E15.5 while blooming for OHC occurs around E16.5. Kinocilium relocalisation reaches its most eccentric position at E16.5 and E18.5 for IHC and OHC, respectively (Figure 3G). We therefore decided to measure HCmv width along the apical-base axis at E16.5 focusing on microvilli in proximity to the kinocilium. We hypothesised that at this time point, we would find the largest heterogeneity along the apical-basal axis. We find that HCmv on the IHCs progressively become wider (from 108±10 nm to 119±16 nm, P=3·10^-4^) while the thickness of microvilli found on the supporting cells and GER cells remain constant (75±10 nm and 72±12 nm, respectively) (Figure 4F). In contrast, the HCmv on OHC do not display any change along the apical-basal axis of OC at E16.5, nor have they thickened significantly (91±15 nm, P=0.06). Altogether, the data suggests that heterogeneity amongst the microvilli of the sensory patch is first observed once hair cells attain their identity and the heterogeneity increases once the kinocilia has radially localised.

We next focused on HCmv width at E18.5, when kinocilium relocalisation had been completed (Figure 3G). At this time point we found a clear distinction between HCmv thickness depending on their proximity to the kinocilium (Figure 4 B-C). The population of HCmv close to the kinocilia are wider (OHC P=2·10^-4^, IHC P<10^-4^) than those further away and the difference in thickness is larger for IHC (146±24 nm versus 102±17 nm) compared to OHC (100±13 nm versus 82±8 nm) (Figure 4G). The entire HCmv population from IHC have increased significantly in width over the course of 3 days (E15.5-E18.5) whereas for the OHC only a subpopulation, those closest to the kinocilium, are thicker than microvilli from support cells of GER cells. This suggests that a subset of HCmv widen into an intermediate state, that we term proto-stereocilia, before they reach their final mature shape.

### Early Post-Natal Development of Stereocilia

Proto-stereocilia widen even more and organise toward their stereotypical row-wise staircase pattern during postnatal development of both IHC and OHC (Figure 5A-B). Proto-stereocilia, closest to the kinocilium at E18.5 (134±22 nm), continue to widen by 38±1 nm per day and can be clearly designated as the first row by P2 and reach a width of 673±38 nm at P12 (Figure 5C). At P2 the second and third row of stereocilia can also be distinguished. Although starting out slightly wider at 282±30 nm (compared to 232±25 nm of the first row at P2), the second row of IHC follows a similar thickening rate as the first (39±2 nm per day), reaching 661±33 nm at P12. In contrast, the third row starts at 148±22 nm, widening more slowly (8±2 nm per day) to reach 223±25 nm at P12. As with IHC, row identity for OHC (proto-)stereocilia becomes evident in the postnatal stage. At P2, rows 1, 2 and 3 attain widths of 152±14 nm, 174±24 nm, and 124±10 nm, respectively (Figure 5D). The width increases slower than on IHC and is similar for all rows being 18±2 nm, 11±4 nm, and 12±1 nm per day, for rows 1,2, and 3, respectively.

Another feature of HC maturation is the resorption of excess HCmv (Kaltenbach et al., 1994). Soon after the appearance of wider HCmv (which we consider to be proto-stereocilia), HCmv distant from the kinocilia begin to be resorbed. However, the rate of resorption is different for IHC and OHC. At E18.5 the HCmv population in the IHC starts falling at a constant exponential rate of 0.21±0.02 day^-1^ (Figure 5E). Although the resorption of HCmv on OHC starts later, at around P0, resorption is faster than on IHC, at 0.40±0.03 day^-1^ (Figure 5F). The difference between the two hair cell types is stark: the IHC loses about half of its microvilli plumage in 3.4 days while the OHC has lost it within 1.7 days. Moreover, the microvilli resorption on OHCs will continue until almost no HCmv are visible post P7. In contrast, around 33±5 HCmv can still be found on IHC at P12.

At P0 the population of the hair cell microvilli that can be assigned as protostereocilia are 59±9 for IHC and 106±12 for OHC. Upon row assignment at P2 these are equally distributed amongst the rows: 20±2, 19±2, and 20±2 for IHC and 33±4, 33±4, and 32±3 for OHCs counting for both rows 1, 2, and 3, respectively. Over time at P12 15-25% of the stereocilia on both IHC and OHC have been resorbed (Figure 5E-F).

Altogether, the data suggests that as the stereocilia mature the hair cell actin machinery simultaneously operates two distinct processes. One designated to resorb and remove actin protrusions and a second tailored to widen and elongate a population of actin protrusions. The commitment of an actin protrusion to one of the processes is likely thresholded by both thickness as well as position.

## DISCUSSION

The development of stereocilia from microvilli is a multi-step process, each guided by different actin-modifying proteins, bundling proteins and motors. These steps generate a precise architecture exquisitely suited to a mechanosensitive role. However, the patterns of actin modifications have been unclear. We employ a new analytical pipeline which uses confocal fluorescence images, to determine signatures of F-actin distribution as HC develop. The advantage of this method is that actin intensities are normalized to cells of the GER. This allows uniform and normalized actin intensities to be extracted from multiple samples prepared from different labs, irrespective of the protocol. Even before any overt morphological distinction, we find that HC can be characterised by higher junctional actin intensities when compared to cortical actin. As HC develop the amount of cortical actin surpasses junctional actin, indicative of the maturation of stereocilia.

Classical work has defined milestones during the development of the hair bundle in inner ear hair cells (Kaltenbach et al., 1994; Tilney et al., 1992b; Zine and Romand, 1996). Prior to HC and SC differentiation, the epithelium consists of uniform cells with sparse microvilli. In the rat, Zine and Romand noted that prior to HC differentiation, two epithelial domains with different microvillar densities, a region abneurally consisted of microvillar sparse cells, and those located neurally were microvillar-rich (Zine and Romand, 1996). Our work in the mouse corroborates this: Two domains based on microvilli density can be observed until E14.5, before the onset of any overt HC or SC differentiation. The epithelium towards the neural edge, which will give rise to the sensory patch and GER, is microvillar-rich. More laterally, towards the abneural edge, the epithelium is microvillar-sparse. We find that one of the first signs of HC formation is the accumulation of junctional actin, although from SEM we did not observe any morphological changes at this stage. However, slightly later, the kinocilium radially relocates, the HC domes, and its microvilli bloom. It is tempting to speculate that the accumulation of the junctional actin is necessary for doming, which itself may be important for the burst of microvilli seen on the HC surface.

Doming has been observed in the endodermal epithelia cells of the rat visceral yolk sac where doming of cell surface correlates with an increase in the number and length of microvilli (Jollie, 1986). Similarly, uterine epithelial cells also exhibit doming with an increase in the microvilli density (Laird et al., 2020). In contrast, enterocytes domes when due to a lack of adhesion molecules that normally support microvilli (Pinette et al., 2019). Indeed, in enterocytes, apical markers are decreased, not increased, as a result of doming. Here, the perturbation of the actin network in brush border enterocytes results in doming and a decrease in microvillar density (Postema et al., 2019). Thus, it is possible that doming represents a state of high actin dynamics.

The gradient of development of hair cells is mirrored in the maturation of the hair bundle (Chen and Segil, 1999; Li and Ruben, 1979; Lim and Anniko, 1985). From the SEM images we are able to capture HC in which the kinocilium has not fully relocalised eccentrically. This is the first stage at which microvilli heterogeneity is observed. Even during kinocilial relocalisation, some of the microvilli are thicker than those found on SC and GER cells, however blooming is only observed once the kinocilium has fully relocalised. As suggested by Tilney and supported here, blooming is associated with the emergence of hexagonal packing of the HCmv (Tilney et al., 1992a; Tilney and DeRosier, 1986). As blooming is only observed in HC in which the kinocilium is eccentric, this may suggest the coupling of these processes. Strong evidence points to the intrinsic polarity pathway, comprising of mInc, LGN and Gαi, as responsible for kinocilial relocalisation (Ezan et al., 2013; Tarchini et al., 2013), however its impact on the kinds of actin dynamics necessary for blooming, is not known.

The next step in the morphogenesis is the elongation of hair cell microvilli closest to the kinocilium. We observe that there is also an increase in width during late embryogenesis, and is particularly prominent in IHC, but can also be quantified in OHC. The coupling of elongation to widening is not clear, however analysis of mutants provides some insights (Krey et al., 2020; Krey et al., 2023). Mutants of Gαi, and Whirlin all show defects in elongation and also show stereocilia that are thicker than in wild types (Manor et al., 2011; Tadenev et al., 2019). In contrast, mutants of Myo15a show a final bundle with elongation deficits and thinner stereocilia (Hadi et al., 2020). However, we note that in these mutants, HCmv are still thicker than surrounding microvilli. Our data indicates that there is a second post-natal thickening step that is necessary for stereocilia to adopt their final form. We speculate that this step, a transition from proto-stereocilia to stereocilia, does not progress in Myo15a mutants. Myosin IIIa and III B are also expressed in stereocilia at embryonic stages of development. Mutations in these proteins perturbs the formation of graded stereocilia and un-resorbed hair cell microvilli (Ebrahim et al., 2016; Lelli et al., 2016). Widening is also perturbed in these mutants, again pointing to a link between elongation and the increase in stereocilium diameter.

The changes in microvilli form as HC mature depend on actin dynamics. By using UN-RaPA to determine actin signatures, we can quantify how mutations in modifiers of actin impact actin dynamics. The normalisation to GER cells has the advantage that any imaging protocol can give a map of actin in the HC. Indeed, UN-RaPA can be used in combination with antibody staining to identify intersectional signatures of a protein with actin. A recent advance, HCAT, uses artificial intelligence and machine learning to identify and segment hair cells in the organ of Corti (Buswinka et al., 2023). By combining this with UN-RaPA to analyse the F-actin and other protein signatures, a powerful analytical pipeline can be developed for phenotyping across many samples, stained at different times in different labs.

## ACKNOWLEDGEMENTS

We are grateful to intramural funding from DAE/TIFR to NCBS, and grants from TIFR Infosys-Leading Edge Grant and Royal National Institute for Deaf People for this research. TSvZ acknowledges a fellowship from the Fundación General CSIC’s ComFuturo programme which has received funding from the European Union’s Horizon 2020 research and innovation programme under the Marie Skłodowska-Curie grant agreement No. 101034263. We would like to thank the Animal Care Resources and Centre at NCBS, The Central Imaging and Flow Facility, the Electron Microscopy facility and lab support at NCBS. We thank lab members for their valuable feedback.

